# Nitrogen response and growth trajectory of sorghum CRISPR-Cas9 mutants using high-throughput phenotyping

**DOI:** 10.1101/2024.12.13.624727

**Authors:** Hongyu Jin, Alexa Park, Avinash Sreedasyam, Guangyong Li, Yufeng Ge, Kankshita Swaminathan, Jeremy Schmutz, Thomas E. Clemente, James C. Schnable, Jinliang Yang

## Abstract

Inorganic nitrogen (N) fertilizer has emerged as one of the key factors driving increased crop yields in the past several decades; however, the overuse of chemical N fertilizer has led to severe ecological and environmental burdens. Understanding how crops respond to N fertilizer has become a central topic in plant science and plant genetics, with the ultimate goal of enhancing N use efficiency (NUE) in crop production. As one of the most essential macronutrients, N significantly influences crop performance across different developmental stages of plant, phenotypic traits result from the accumulative effects of genetic factors, prevailing environmental conditions (specifically N availability), and their complex interactions. To characterize the targeting N-responsiveness and growth trajectory, we employed CRISPR-Cas9 technique to generate sorghum mutants using CRISPR technology. Using a LemnaTec plant imaging system, we obtained time series imagery data from 29 to 130 days after sowing (DAS) for these CRISPR-edited mutants under high N and low N greenhouse conditions. After imagery data analysis, we extracted a number of morphological and greenness index traits as a proxy of plant growth and N responses. Subsequently, we employed two different methods to model the temporal N-responsive traits, allowing us to estimate seven key parameters from the growth curve. Our findings revealed that the wildtype and the edited sorghum lines exhibited differences in N responses for several of the key growth-related parameters. The high-throughput N phenotyping pipeline paves the way for a better understanding of the N responses of edited lines in a dynamic manner and sheds light on further improvements in crop NUE.

## Introduction

Modern agriculture has become increasingly mechanized and specialized, heavily reliant on fossil fuels as the primary energy source and on the application of large quantities of chemical fertilizers to improve productivity. Since the invention of the Haber–Bosch process, nitrogen (N) fertilizer has been increasingly used in crop production, leading to 30% to 50% of crop yield gain (1; 2). Global data for the use of N fertilizer for the three major cereal crops (maize, rice, and wheat) indicate that only 18% to 49% of the applied N is taken up by these crops, with the remainder being lost to runoff, leaching into waterways, and volatilization (3; 4). The excessive application of inorganic N fertilizer has adverse effects on biodiversity and poses threats to human and animal health within the agroecosystem (4). The increasing demand for sustainable agriculture provides new challenges for plant breeding and genetics, where achieving comparable yields while maintaining soil and environmental health requires reducing chemical fertilizer inputs and minimizing carbon footprints (5). In addition, CRISPR-Cas9-based gene editing has provided unprecedented efficiency in mutant genesis. Therefore, it is critical to investigate the genes responsible for crop N responses via reverse genetics to inform future plant breeding practices.

Sorghum is the fifth most-grown cereal crop globally, with approximately half a billion people, mainly in developing countries, relying on it for food security (6). In contrast to recently developed maize hybrids optimized for high N cultivation to enhance economic productivity, such as in the U.S. Corn Belt (7; 8), sorghum cultivars exhibit significant diversity in N use efficiency (NUE), stemming from their natural adaptation in varied fertility environments (9). The NUE diversity positions sorghum as an ideal model C4 crop for investigating plant responses to N deficiency and NUE in crop production. N deficiency has been reported to systematically impact various aspects of sorghum growth, spanning from morphological traits, such as leaf area, tiller number, and plant height, to developmental characteristics, like flowering time and leaf senescence, as well as physiological traits, including chlorophyll content, protein content, plant hormone metabolism, and membrane transport, among others (10–13). These changes in phenotypes ultimately reflect the interactions between the plant genetics and its low N environment, leading to decreased photosynthesis efficiency and reduced biomass production (14; 15).

To quantify plant N response, traditionally, researchers have used laboratory-based biochemistry methods to measure plant N and chlorophyll content, and manually collected plant morphological traits (10). These methods are relatively cost- and labor-intensive, which often limits the sample density and sample size. High-throughput phenotyping is a set of phenotyping techniques, primarily focused on automated data acquisition at a large scale, that has recently emerged as an enabling tool for crop breeding and plant science research. Numerous high-throughput phenotyping studies have employed spectral or imaging techniques using sensors or cameras. These techniques have captured information ranging from red-green-blue (RGB) color to infrared, thermal, multispectral, and hyperspectral data (16–21). Previous studies have demonstrated that plant structural phenotypes obtained from high-throughput phenotyping experiments, such as plant height, convex hull area, and pixel count, can be used as indicators of N levels in plants (22; 23).

Using a camera-based high-throughput phenotyping system coupled with automation, massive amounts of imagery data can be obtained daily, presenting new challenges for image data analysis and result interpretation. High-throughput phenotyping captures information about plants and their environment with inherent noise. Domain knowledge in plant biology and appropriate experimental design are essential for developing models that distinguish noise from informative variables, which may mediate, covary with, or interact with the features of interest. Additionally, high-throughput phenotyping enables non-destructive measurements across crop growth stages and often generates high-dimensional temporal data. An intuitive and widely applicable approach is to treat each single time point as a phenotype. Comparisons between the factors of interest can then be directly made for each time point by statistical models such as ANOVA, after which the biological significance is summarized (24; 25). To gain deeper insight into the mechanisms underlying crop development, functional analysis of variance (FANOVA) was employed to decompose the temporal variance into factors such as genotype and treatment (26; 27). Functional principal component (FPC) analysis, a functional-based factor model, has been shown to associate FPCs representing growth trajectory characteristics with genomic markers or QTLs (28; 29). Recently, researchers have used latent space phenotyping methods to automatically learn hidden parameters that represent the response to treatment and are associated with the genomic data (30–32). These approaches captured overall visually evident features from plant imagery data or spectra data without the need to construct and extract features using conventional modeling methods. However, the interpretation of such latent space phenotypes can be challenging.

In this study, leveraging the in-house generated CRISPR-Cas9 mutants in sorghum, we present a pipeline for characterizing plant growth responses to different N treatments. The pipeline utilizes the automated plant imaging system, combining it with computer vision methods to extract phenotypes and statistical modeling to summarize the high-dimensional temporal phenotype into biologically meaningful parameters. This data acquisition and analytical pipeline paves the way for a better understanding of N responses in CRISPR-Cas9 edited mutants, shedding light on facilitating plant breeding.

## Results

### A high throughput phenotyping procedure to characterize sorghum mutants for N responses

In oura previous study (33), researchers we performed a comparative transcriptome analysis to investigate nitrogen response across various plants. Specifically, sorghum’s response was examined to varying nutrient availability (ammonium, nitrate, and urea) by analyzing both differential gene expression and co-expression patterns. A nitrogen-responsive co-expression module in root tissue was identified, centered around two hub genes: Sobic.001G040200 (encoding a NAC transcription factor) and Sobic.002G121300 (encoding a serine carboxypeptidase). Further analysis of the immediate network of these hub genes focused on their direct nodes to assess their potential roles in nitrogen metabolism and assimilation. This investigation led to the identification of 12 genes within the module with putative nitrogen-related functions. In the current study, we worked on in-depth characterization of six of these 12 genes (**Table S1** and **S2** and **Figure ??**) Using CRISPR-Cas9 technology (34), we obtained five gene-edited sorghum lines in Tx430 background, with up to three genes being knocked out for each homozygous mutant (see **Table S1** and **S2**). These gene-edited sorghum mutants and Tx430 wildtype (WT) were grown at the LemnaTec high-throughput plant phenotyping platform with two different N treatments: high N (15 mM nitrate) and low N (3 mM nitrate) (35). The experiment was conducted following a complete randomized block design with three replications per genotype (**Figure 1A, Materials and Methods**).

**Figure 1.**
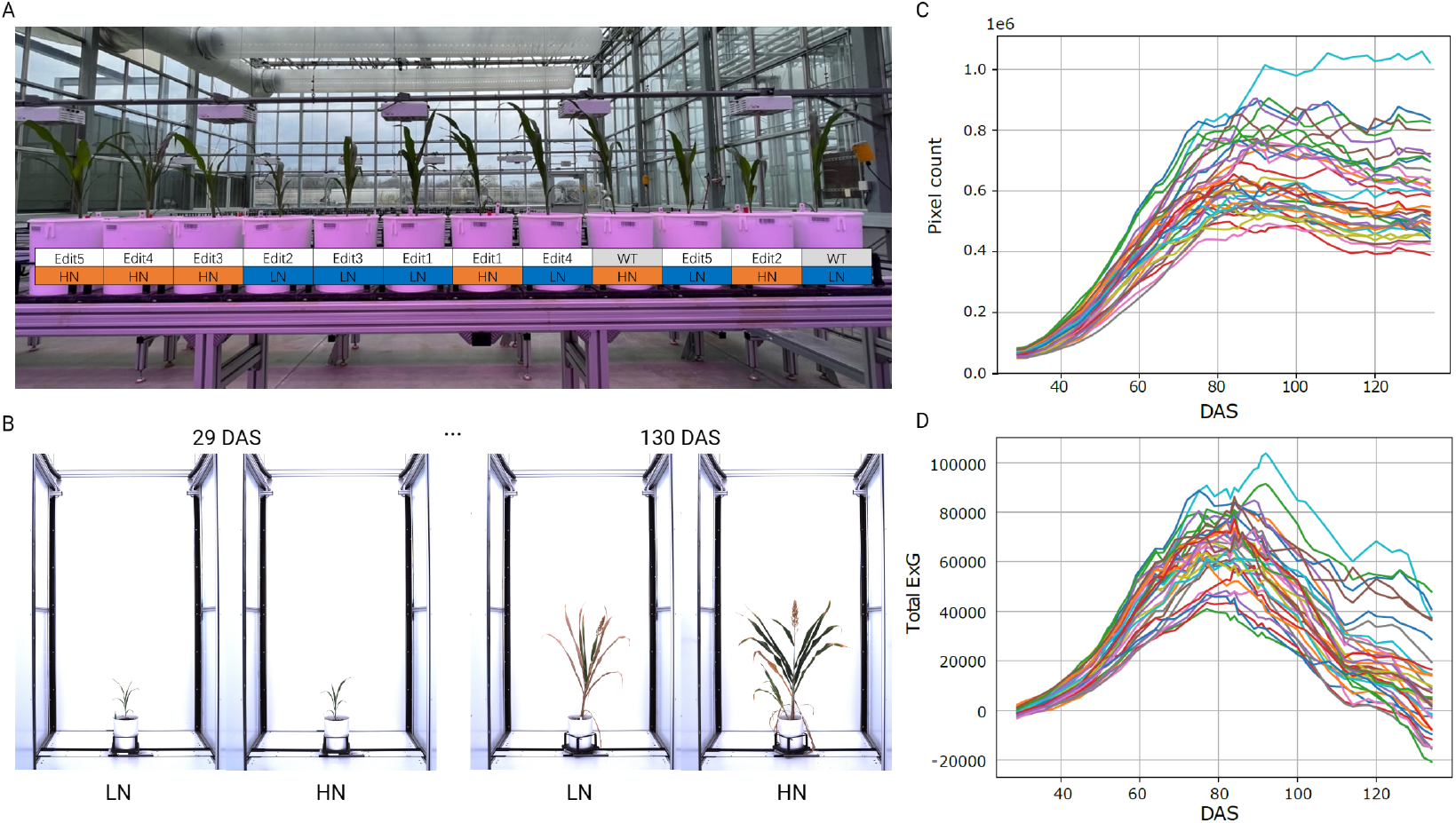
Characterizing sorghum CRISPR mutants for N responses using a high throughput phenotyping procedure. (**A**) The greenhouse experimental design for CRISPR mutants under high N (15 mM) and low N (3 mM) conditions. (**B**) Time-series automatic image acquisition from the first day of phenotyping (29 DAS) to the last day of phenotyping (130 DAS) The examples are Tx430 WT. The extracted phenotypes (**C**) pixel count and (**D**) total ExG index value for each individual during the phenotyping period.

We also conducted conventional phenotyping for plant dry weight at the endpoint of the experiment. The Tx430 WT showed significantly increased dry weight accumulation under high N conditions in each part of the plant we measured, indicating its sensitivity and response to N treatment. The same trend was found for Edit4 and Edit5. Edit1, however, exhibited no significant difference in any of the dry weights between the two N treatments (**Figure S1**). Edit2 and Edit3 had moderate phenotypes that only the whole plant dry weight showed significant N responses (F-test, p-values < 0.05). These conventional phenotyping results suggested that N treatments significantly influenced sorghum growth, with distinct responses observed among various gene-edited lines. Tracking N responses across specific growth stages could reveal important details about how the N responsiveness has changed in these gene-edited lines.

For the high throughput phenotyping, in total, 15,120 imagery data points were obtained in a time series manner at 42 different time points (**Figure 1B**). We used a manually created background image for background subtraction, followed by image opening and closing morphological operations to remove the potential “salt and pepper noise” that remained in the background (36). This ensured more successful segmentation across different growth stages (**Figure S2**, see **Materials and Methods**), comparing to color thresholding, where green tissues, panicles, and senescent tissues require different thresholds. Two phenotypic traits associated with plant growth and N content were extracted in a time-series manner, from which growth trajectories were then modeled and explained (**Figure 1C and D** and **Table S1 and S2**).

### Parameterization of pixel count growth trajectories

The plant pixel count has been previously reported to be strongly correlated with fresh weight, which serves as a proxy for biomass (37). To study biomass dynamics using our time-series imagery data, pixel counts for each plant on each imaging date were extracted from the segmented images and averaged for 10 different angles of side-view images. The plant pixel counts in our experiment followed an S-shaped curve (**Figure 2B**), which can be fitted as a logistic function, consistent with a number of previous studies (29; 38). We estimated the steepness, inflection point and amplitude parameters (*k, µ*, and *A*) from the logistic growth curve by fitting the model (see **Materials and Methods** and **Figure 2B**).

**Figure 2.**
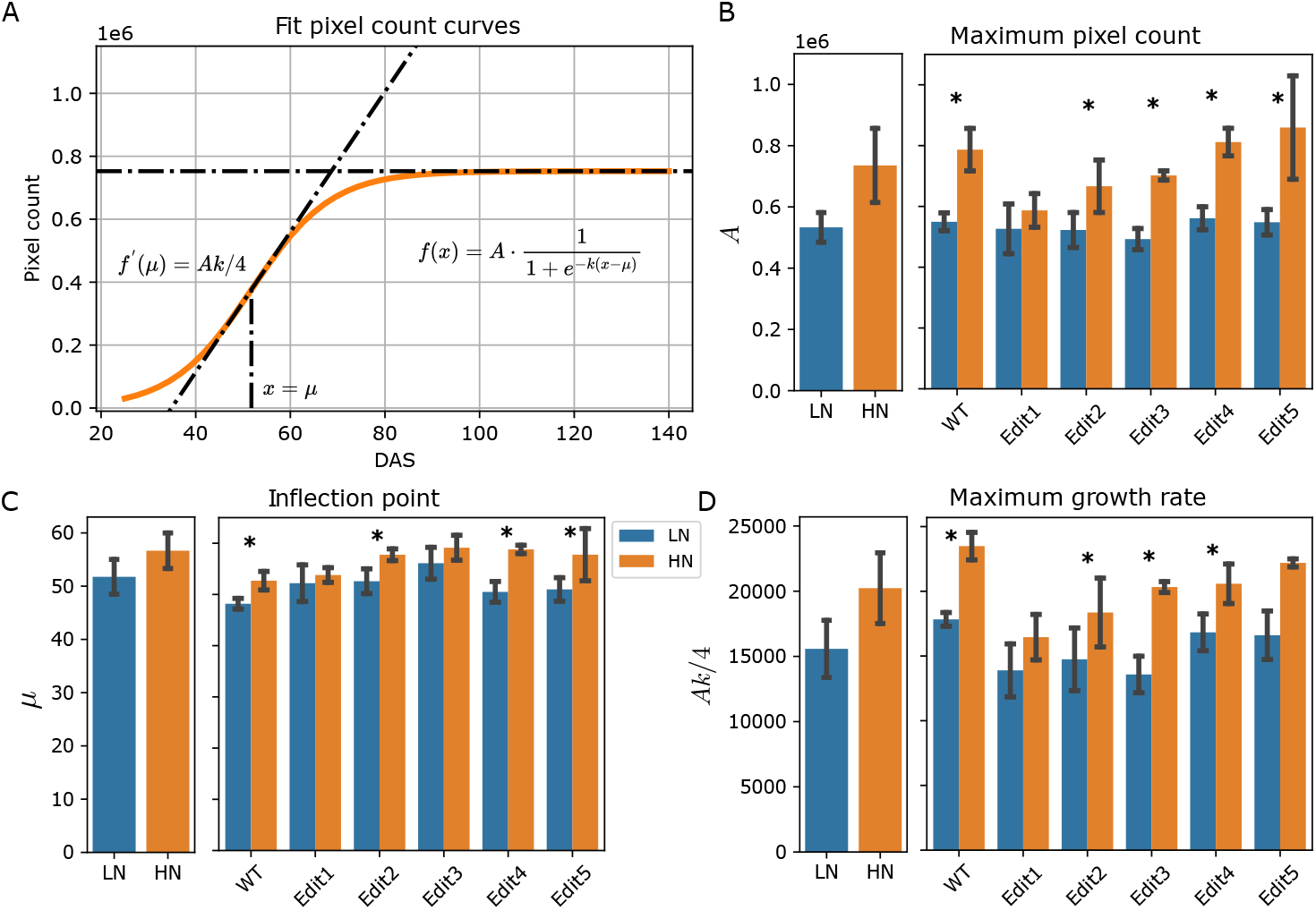
Modeling of the pixel count trait. (A) Logistic curves with the parameters estimated from the data. The bold lines represent the curves of WT Tx430 plants, and the dashed color lines represent the gene-edited lines. (B) (C) and (D) The phenotypes estimated or derived from the function. The left panel shows the overall comparisons between treatment and the right panel shows the comparisons for each genotype. The error bars indicate the standard deviation. Stars indicate significant difference between low N and high N treatment for the pooled data or within a single genotype.

After extracting estimated parameters, the results suggested that the maximum pixel counts (*A*), serving as a proxy for total biomass, exhibited significant (ANOVA, p-value < 0.050) genotype effect and treatment effect (**Figure S3** and **Table S3**). When compared with the low N treatment group, overall significantly (F-test, p-value < 0.001) higher maximum pixel counts under high N condition was observed (**Figure 2B and Table S3**). Particularly, Tx430 WT and Edit3-5 showed significantly (F-test, p-value < 0.050, **Table S3**) better biomass (or *A* value) in high N than low N but the trend was not found in Edit1 and Edit2. This observation highly correlated with the dry weight measurements (**Figure S1**).

The parameter *µ*, representing the DAS when the fastest growth rate was achieved, suggested that overall the plants achieved the fastest growth rate later under high N than low N conditions (F-test, p-value < 0.001, **Figure 2C** and **Table S1**). Again, this pattern was not observed for Edit1 (F-test, p-value= 0.312, **Table S3**). Consistently, the fastest growth rate, measured using *Ak/*4 (see **Materials and Methods**), indicated that there was overall significant faster rate (F-test, p-value < 0.050, **Figure 2D** and **Table S3**) when N was sufficient. Taken together, the data were consistent with our observation that WT and Edit2-5 grew longer and faster, accumulating more biomass under high N compared to low N conditions, while Edit1 appeared unresponsive to high N treatments.

### Characterization of phenotypic dynamics using Excess Green indices

We next sought to model a different trait, Excess Green (ExG), extracted from the segmented images. ExG is a RGB color-based vegetation index that has been used to describe the greenness of vegetation, indicating the N content (25; 39–41). The ExG increased at the beginning of the experiment and started to decrease on around 80 DAS. Ten individuals showed total ExG below 0 at the end of the experiment. This might indicate that the plants were largely senescent and the ExG value of the yellow-colored pixels was negative. A recent study has shown that the bell-shaped Normalized Green-Red Difference Index (NGRDI) growth trajectories can be modeled by a Gaussian peak function (29). Similarly, we used a Gaussian function with an additional *y* location parameter to model our total ExG values for each individual (**Figure 1D, Figure 3A**, and **Table S2**).

**Figure 3.**
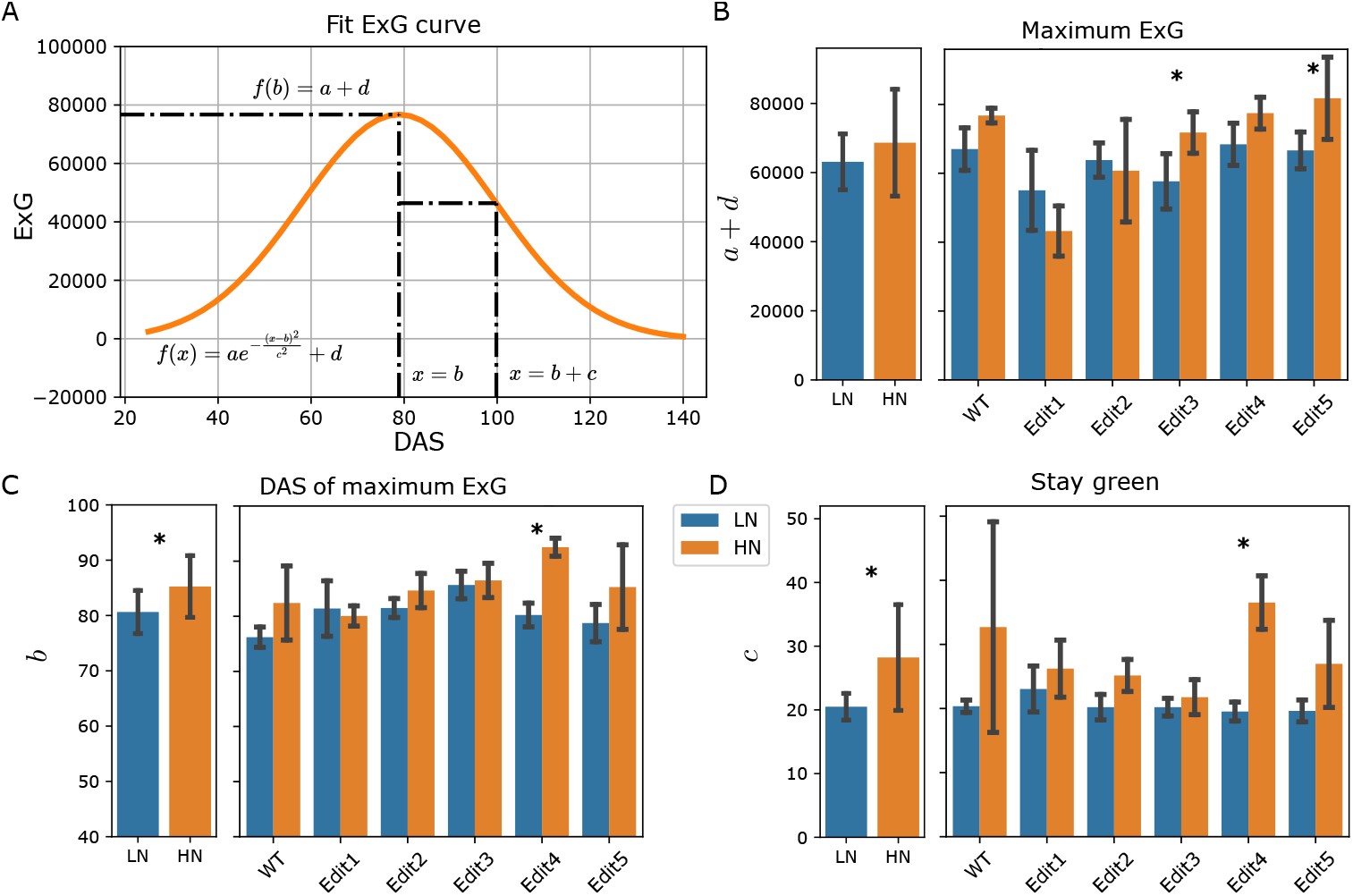
Modeling of the ExG index. (A) Fitting the ExG curve as Gaussian curve and extraction of meaningful phenotypes. (B)-(D) The comparisons of the extracted phenotypes between N treatments. The left panel shows the overall comparisons between treatment and the right panel shows the comparisons for each genotype. The error bars indicate the standard deviation. Stars indicate significant difference between low N and high N treatment for the pooled data or within a single genotype.

The maximum ExG, revealed by the combination of *a* + *d* (see **Materials and Methods**), showed a significant reduction in Edit1 and Edit2 compared to Tx430 WT under high N condition (F-test, p-values < 0.05, **Figure S4**). When considering low N treatment, there was a marginally significant N treatment effect for the maximum ExG (**Figure 3B** and **Table S2**). Specifically, Edit3 and Edit5 showed significant N responses (F-test, p-value = 0.045 and 0.033, **Table S4**) between the two treatments.

The DAS achieved the maximum ExG was revealed by the *b* parameter. The high N condition led to an overall significant (F-test, p-value= 0.016) later DAS to achieve the maximum ExG (**Figure 3C** and **Table S4**). The *c* parameter indicated the time between the DAS of the maximum ExG and the DAS of the maximum ExG growth or decay, which we defined as a measurement of stay green phenotype (see **Materials and Methods**). A larger *c* parameter indicates the plant may stay greener for a longer time. Plants under high N generally stayed green for longer (F-test, p-value < 0.001). Per-genotype level significance for these two parameters was only observed in Edit4 (F-test, p-value = 0.001, **Figure 3D**). Together with the maximum ExG and DAS of maximum ExG traits introduced above, these may suggest sufficient N prolongs the plant’s vegetation while N deficiency might stimulate plant to complete its reproductive cycle before resources were depleted.

## Discussion

In this study, we phenotyped 5 gene edited sorghum lines in high N and low N environment in comparisons to the Tx430 WT control. Tx430 WT demonstrated a growth-optimized strategy under high H compared to low N. It enriched chlorophyll content and accumulated the most biomass by taking full advantage of N resources. The edited lines showed compromised strategies with limited ability to capitalize on high N but significantly less than the WT. Edit1, especially with three N-related genes knocked out simultaneously, appeared to default to a stress-conservation strategy, even when N resources were abundant. This could occur due to disrupted sensing or signaling pathways that fail to perceive high N as a growth-promoting cue, but further investigations for the pathways that each of the genes are involved in are needed.

This study introduced a new high throughput phenotyping framework that discovered and quantified sorghum N responsive traits. By estimating the parameters from the time-series growth curves, we were able to extract not only the conventional end-point traits, such as above ground biomass and N content, but also growth trajectory-related phenotypes, such as critical timings. In this study, our models’ parameters identified or derived several important dates or stages after sowing. The treatments and phenotyping started on 29 DAS, on which the plants from both treatment groups were at the five leaf stage (**Figure 1**). The first key time point we modeled was the inflection point (*µ*), representing the DAS at which the plant achieved the highest growth rate of the pixel count. The Tx430 WT grown in low N condition was at the eight leaf stage (**Figure S5**). Following the inflection point, the next time point was the critical point (*b*) of ExG, the time that reached the highest total ExG. The overall later time of reaching the highest ExG in high N indicates the N effect that prolonged the increasing of greenness or potentially the photosynthesis intensity. The third time point our models identified as important is the time of the maximum decay of ExG (*b* + *c*). This may provide insights of the stay-green phenotype in response to N condition. These are important in N response because in response to abiotic stresses, plants usually exhibit different strategies to balance the growth and stress response trade-off. Some plants may accelerate their maturity as a form of stress escape, while some may delay their maturity to accumulate the resources for successful reproduction (42–44). By looking at the growth and senescence rate, and the specific timing (such as inflection points in this study) in growth stages, we are able to notice the difference of the sorghum plants in response to N treatments. Moreover, identifying different growth trajectories improves the comparability of the two end-point traits since traditionally researchers usually select a specific date to measure all the plants, which results may be confounded with the growth stage.

Sustainable crop production is imperative to ensure food security for both present and future generations. Enhancing N use efficiency in plants is crucial for achieving environmental and ecological sustainability. Understanding the N deficiency phenotype is fundamental for breeding crops with improved N utilization. Traditionally, biochemical methods (e.g., chlorophyll or N content analysis) were employed to assess N deficiency (45). Alternatively, the plant size and color can be used as an indicator, but quantifying with human’s eyes may lead to low precision. With the fast development of image-based high throughput phenotyping, our pipeline quantified the sorghum plant color and size in different N conditions over the growth stages. To deal with the high dimensional temporal data, we employed mathematical modeling approaches, which summarize the functional data into several biological meaningful parameters, which were also used to derive more qualitative descriptions to the N-related phenotypic traits.

As high-throughput and high-intensity plant phenotyping are developing, time series repeated measurements are becoming increasingly achievable for the researchers. This poses a challenge of developing appropriate models to analyze the phenotypic dynamic of plants. In this study, we summarized high-dimensional time-series traits into N-responsive phenotypic traits using two mathematical models. This approach modeled the plant growth trajectories with available measurements from the experiment. Compared to the approach that analyzes each single time point separately, our approach is more statistically and biologically meaningful to incorporate information from nearby time points. It also reduces the post hoc efforts for interpretation after the single time point comparisons. There are several attempts using non-parametric functional models to address this problem (26–29). Miao et al., (2020) (28) used a functional principal component (FPC) analysis to extract the main variations in the growth curve of the height of the sorghum and used FPC scores as phenotypes to identify genomic markers associated with the height of the plant. This method is effective in distinguishing between genotypes or treatments, but the interpretation of each FPC remains challenging without pre-annotated genetic associated data. Another non-parametric model, FANOVA has also been applied to dissect the variance of genotypes and treatments (26) but the resulting comparisons are represented as functional data and may present challenge to understand. On the contrary, the mathematical modeling approach reduced the dimension of the time-series data to well-defined parameters, which can be directly linked to biological process. It is also less prone to the hyperparameters of the model and the random noise in the growth curve. In addition, the resulted mathematical models are effective for predicting the unobserved phenotype of the plants. Along with the widely used sigmoid function for plant growth (29; 38; 46), our study identified the ExG growth pattern as a Gaussian peak function with an additional vertical adjustment. This implied that based on the pattern of the growth dynamic, various mathematical functions might be constructed or adjusted to describe the data. Despite of several advantages of modeling the temporal phenotype with mathematical functions, it is worth noting that no model is correct. The mathematical models smooth the data based on functions which require prior assumptions. Important features might be lost if the assumptions misrepresent the data. We noticed that the unbalanced growth and decay rates of the ExG curves can be better captured by skewed Gaussian peak functions, but the additional parameter may introduce more uncertainty and difficulty in the biological interpretation of the parameters. This study presented the feasibility of a high throughput temporal phenotyping pipeline for sorghum N responsive traits. It is promising to be used in future studies but more replication and careful experimental design will be required to boost the statistical power for accurate gene functional characterizations.

## Materials and Methods

### Plant growth and imaging

Five gene-edited sorghum lines (named Edit 1 to 5, **Table 1**) and Tx430 control were planted at Greenhouse Innovation Center of University of Nebraska-Lincoln in small pots on March 23, 2022. Healthily germinated plants were transferred to 1.5 gallon pots on April 14, 2022. Each 1.5 gallon pot contained soil composed of two thirds peat moss, one third vermiculite and 2000 g/yd of lime, which contained no N fertilizer. From planting to April 21, 2022, 20-10-20 N-phosphorus-potassium liquid fertilizer was applied to saturation to ensure the seedlings’ survival.

The plants were placed on the belts of the LemnaTec3D automated imaging system (LemnaTec GmbH, Aachen, Germany) following a randomized complete block design, which contained 3 blocks and each block consisted of 12 plants, a full replication of 6 genotypes grown in 2 N conditions. 53.6 g N-free Hoagland medium powder (bioWORLD Molecular Life Sciences, USA) and potassium nitrate (60.66 g for high N or 12.13 g for low N) was diluted to 40 liters with fertilizer-free water in the greenhouse to make 15 mM N or 3 mM N nutrient solutions. The solutions were applied 250 mL to each pot twice a week. The same solutions were prepared every two weeks and stored in the dark room to avoid the growth of algae Additional water was applied to each pot every day. In total, we imaged 36 plants. The imaging started on April 21, 2022 and terminated on August 4, 2022.

### Dry weight measurement

We harvested the whole plants from the experiment on August 4th, 2022 or 130 DAS. The roots were carefully shaken and washed, after which the plants were dried in a 65 ^*°*^Cdryer for 5 days. Then the dry weight of the plant was measured separately for shoot, root, and panicle.

### Image segmentation

To segment the plant pixels from the background, we used a customized python script with multiple computer vision functions imported from OpenCV 4.1.0.25 (47). More specifically, we constructed a binary mask for the plant by subtracting the pixel values of a background-only image from every plant image, followed by Otsu binarization (48) of the green channel. Ideally, only the plant area has non-zero pixels in the resulting binary mask, but some noises may exist. To denoise, we cropped the mask and the image from the four edges of the image to remove the residue from the frame of the imaging chamber and the pot due to imperfection of overlapping between the plant image and the background image. We applied image opening and closing morphological operations with a 5-by-5 square structure element to remove the salt and pepper noise. Finally, we multiply the binary mask with the raw image to get the segmented image.

### Phenotypic traits extraction from imagery data

We used the segmented images to extract plant phenotypic traits, including pixel-coordinates-based traits such as plant height, plant width and pixel coverage, and color-based vegetation indices. The plant pixel coverage was measured as the total number of non-zero pixels in a segmented image. This number is comparable across the images since all the images are of the same dimension. The plant height was measured from the vertical axis value of the first non-zero pixel from the bottom to the first non-zero pixel from the top. The plant width was measured from the horizontal axis value of the first non-zero pixel from the left to the first non-zero pixel from the right. The formulas of vegetation indices calculation were as shown by Rodene et al (25). Briefly, the values of the RGB channels were used and average values across all the segmented pixels were recorded.

### Statistical modeling of growth dynamics

We used the following logistic function to model the sorghum pixel count growth dynamics. The pixel count on the x^th^ day after sowing is modeled as:

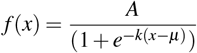

where *A* was the carrying capacity, *µ* was the inflection point and *k* was the steepness. The first derivative of the pixel count:

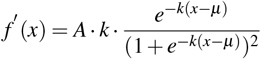

was used to calculate the slope at the inflection point *f* ^*′*^ (*µ*) = *Ak/*4, which represented the maximum growth rate.

We used a Gaussian peak function to model the sorghum ExG index growth dynamic. The ExG on the x^th^ day after sowing is modeled as:

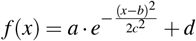

where *a* was the amplitude, *b* was the critical point, *c* was the width parameter and *d* was the vertical adjustment of the curve. The maximum ExG, which was reached when DAS equaled *b*, was controlled by both the amplitude and the vertical adjustment, therefore, it was calculated as *a* + *d*. In addition, the *c* parameter described the width of the peak, which we considered as a measurement of the stay-green phenotype. It is equal to the distance between the DAS of the peak and the inflection point, which shows highest or lowest slope, at either side.

All the parameters were solved for each of the 36 plants using nonlinear least squares regression as implemented in Scipy v1.12.0 (49). The parameters and derived traits were analyzed by analysis of variance (ANOVA). The differences of high N and low N within each genotype were compared using F-contrasts.

## Acknowledgements

This project was supported by the US Department of Energy grant number DE-SC0023138 and the National Science Foundation under the award number OIA-1826781.

## Author contributions statement

H.J., Y.G., K.S., T.E.C., J.C.S. and J.Y. conceived the experiment(s), H.J., A.P. and G.L. conducted the experiment(s), H.J. and A.S. analyzed the results. All authors reviewed the manuscript.

## Supporting Information

### Supporting Tables

**Table S1**. The phenotypic values calculated from the pixel count curves.

(https://github.com/JIN-HY/Sorghum-edits-N-Phenotyping/blob/main/fitpx.csv)

**Table S2**. The phenotypic values calculated from the ExG curves.

(https://github.com/JIN-HY/Sorghum-edits-N-Phenotyping/blob/main/fitexg.csv)

**Table S3**. The contrasts of the phenotypes calculated from the pixel count curves.

(https://github.com/JIN-HY/Sorghum-edits-N-Phenotyping/blob/main/PXcontrasts.xlsx)

**Table S4**. The contrasts of the phenotypes calculated from the ExG curves.

(https://github.com/JIN-HY/Sorghum-edits-N-Phenotyping/blob/main/ExGcontrast.xlsx)

### Supplementary figures

**Table S1.**
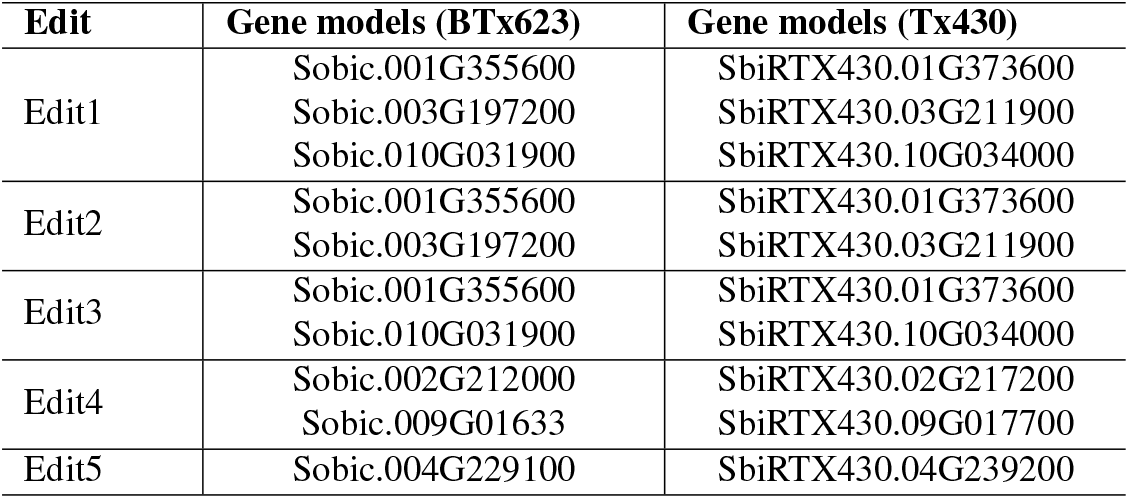
Genes being knocked-out in the 5 edits.

| Edit | Gene models (BTx623) | Gene models (Tx430) |
| --- | --- | --- |
| Edit1 | Sobic.001G355600 | SbiRTX430.01G373600 |
|  | Sobic.003G197200 | SbiRTX430.03G211900 |
|  | Sobic.010G031900 | SbiRTX430.10G034000 |
| Edit2 | Sobic.001G355600 | SbiRTX430.01G373600 |
|  | Sobic.003G197200 | SbiRTX430.03G211900 |
| Edit3 | Sobic.001G355600 | SbiRTX430.01G373600 |
|  | Sobic.010G031900 | SbiRTX430.10G034000 |
| Edit4 | Sobic.002G212000 | SbiRTX430.02G217200 |
|  | Sobic.009G01633 | SbiRTX430.09G017700 |
| Edit5 | Sobic.004G229100 | SbiRTX430.04G239200 |

**Table S2.** Genes targeted for CRISPR-Cas9 editing.

| BTx623 | Tx430 | Description |
| --- | --- | --- |
| Sobic.001G355600 | SbiRTX430.01G373600 | basic helix-loop-helix transcription factor (bHLH47) |
| Sobic.003G197200 | SbiRTX430.03G211900 | 1-aminocyclopropane-1-carboxylate oxidase 1-like protein |
| Sobic.010G031900 | SbiRTX430.10G034000 | early nodulin 93 ENOD93 protein |
| Sobic.002G212000 | SbiRTX430.02G217200 | ethylene-responsive transcription factor 1-like (EREBP) |
| Sobic.009G016633 | SbiRTX430.09G017700 | MYB-like DNA binding transcription factor (MYB59-like) |
| Sobic.004G229100 | SbiRTX430.04G239200 | F-box protein |

**Figure S1.**
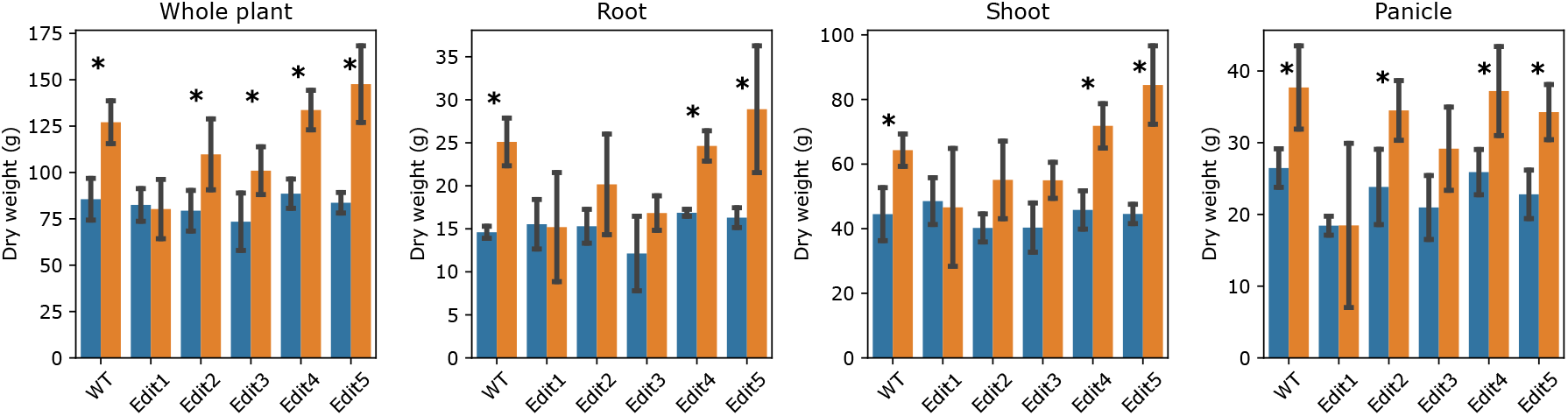
Comparisons of the hand measured dry weight traits. Stars indicate significant difference between low N and high N within the genotype.

**Figure S2.**
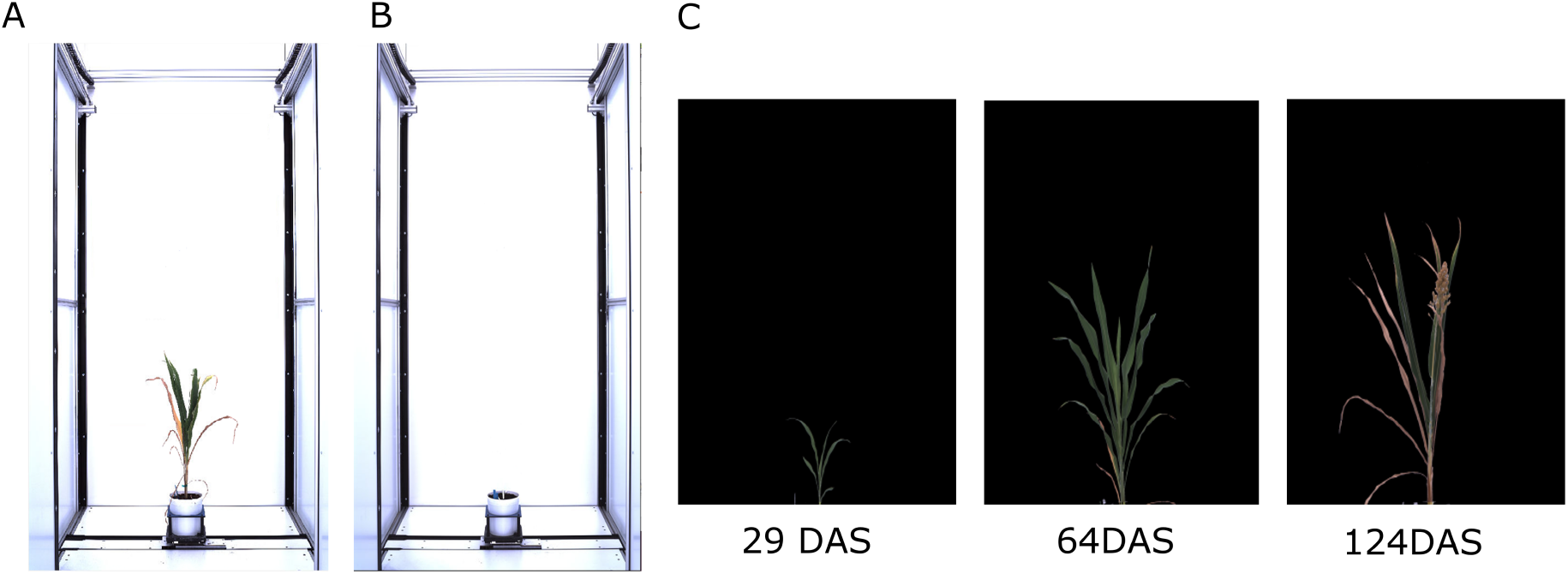
Image segmentation by background subtraction. (A) Example of a raw image. (B) The background image. (C) Segementation results for images taken from different growth stages.

**Figure S3.**
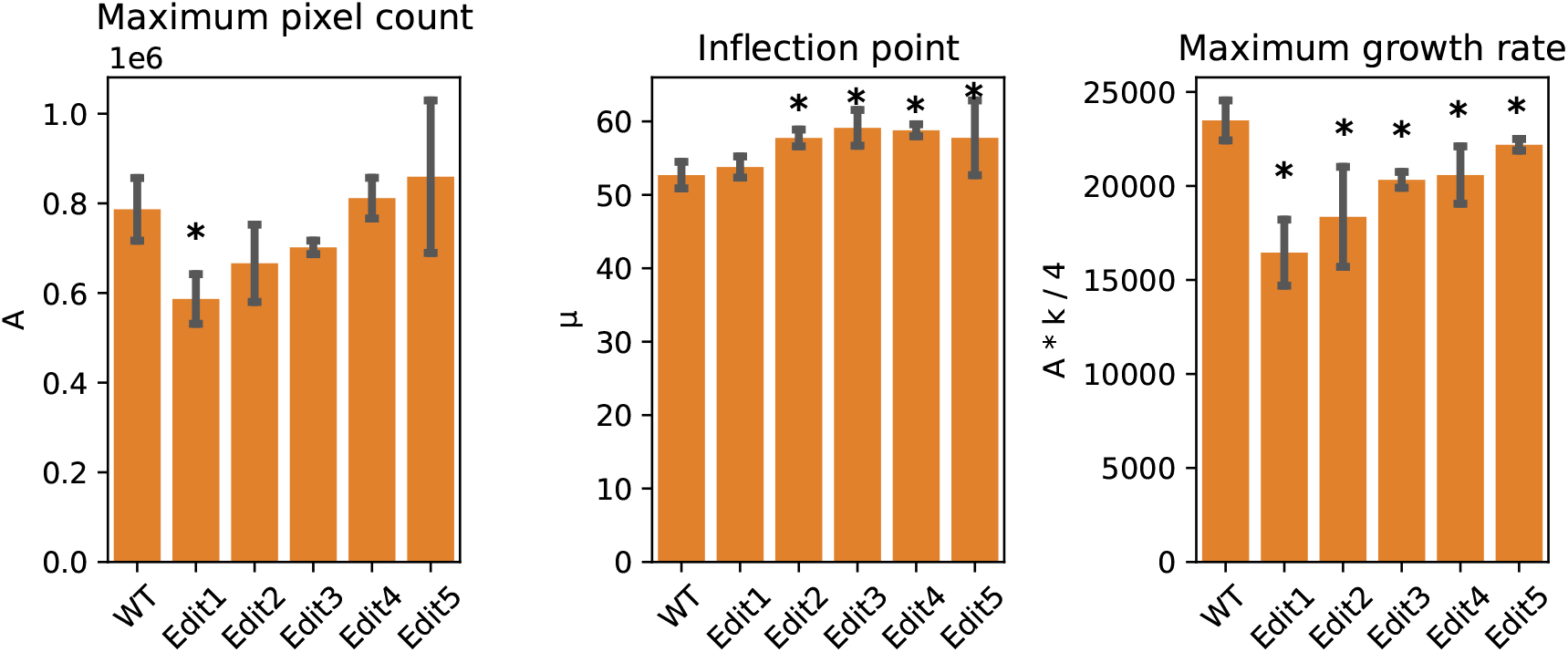
Extracted phenotypes from pixel count curves under normal condition (high N). Stars indicate significant difference between the genotype and Tx430 WT.

**Figure S4.**
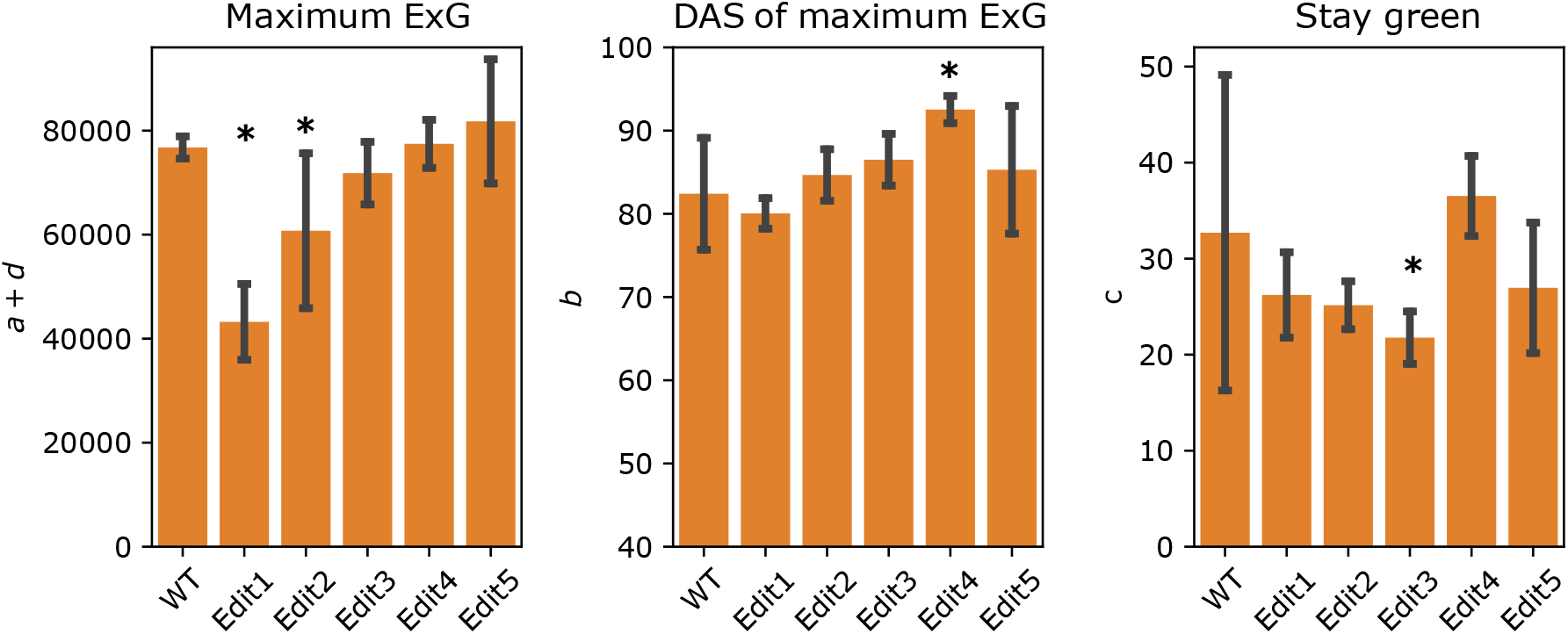
Extracted phenotypes from ExG curves under normal condition (high N). Stars indicate significant difference between the genotype and Tx430 WT.

**Figure S5.**
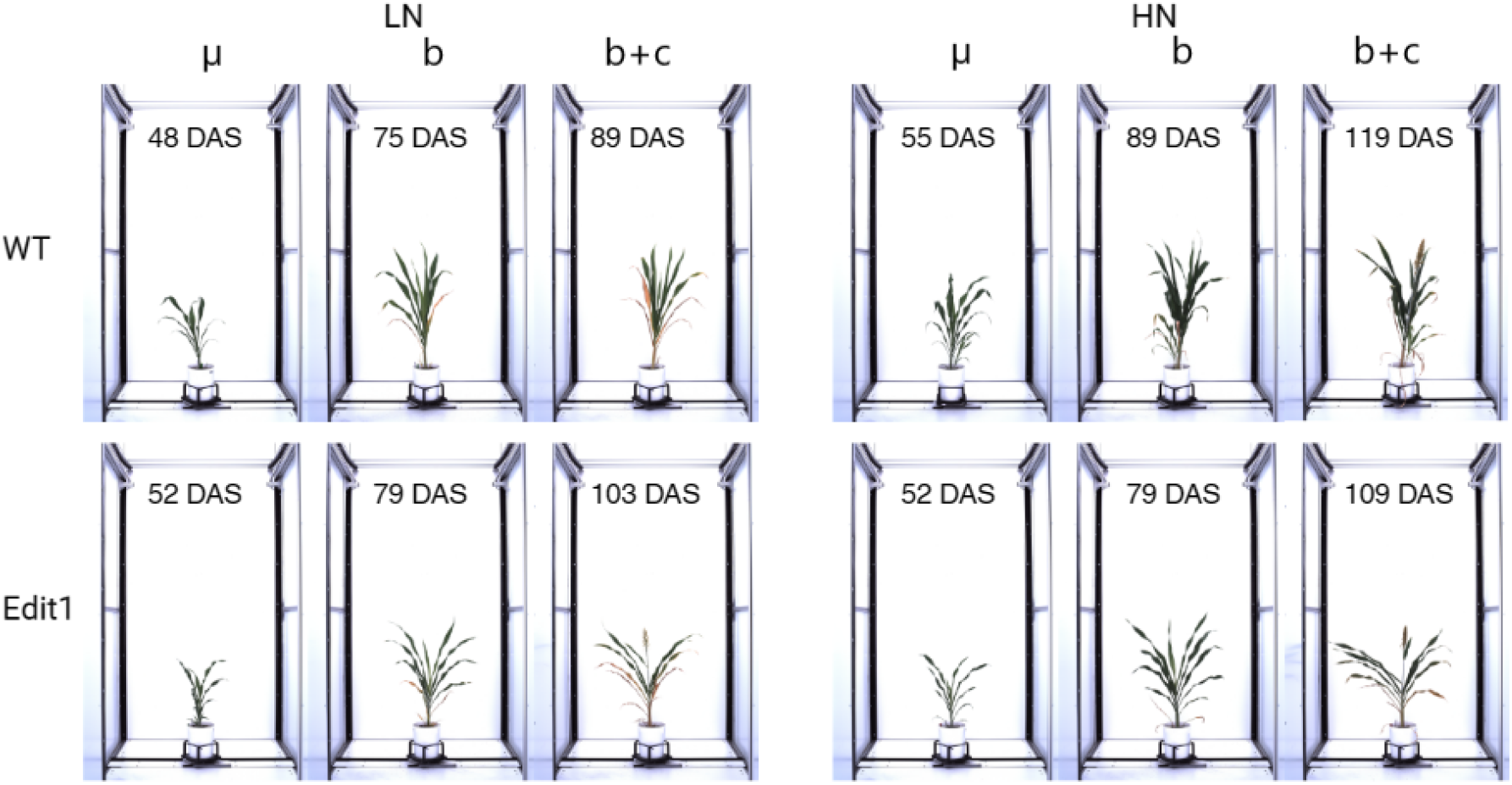
Photos on the date points identified by the models. Photos show the 0 degree side view angle. Tx430 WT and Edit1 from the first replicate were used as the example.

